# Anhedonia buffers the effects of early-life unpredictability on threat-reward decision-making

**DOI:** 10.64898/2026.05.16.725643

**Authors:** Bianca T. Leonard, Maria Alejandra Martinez-Ortiz, Jason Bock, Yifan Zhang, Steve Flores, Derek Vincent Taylor, Laura M. Glynn, Elysia Poggi Davis, Hal S. Stern, Tallie Z. Baram, Catherine A. Hartley, Michael A. Yassa, Aaron Bornstein

## Abstract

Anhedonia – the diminished capacity to experience or anticipate pleasure – is among the most common consequences of early-life unpredictability, yet how these co-occurring conditions jointly shape real-world decision-making remains unknown. Here, we use a sequential foraging-under-threat task to probe motivational conflict decisions in 357 individuals varying in early-life unpredictability and anhedonia symptoms. We find that unpredictability and anhedonia exert *opposing* influences on choice: unpredictability shifts behavior away from the survival-optimal policy in a sex-dependent manner, while anhedonia promotes adherence to it, partly through heightened sensitivity to unexpected threatening outcomes. A mediation analysis reveals that anhedonia partially buffers the deleterious effects of unpredictability on decision quality. These results demonstrate that co-occurring conditions can mask one another’s behavioral signatures and suggest that the heterogeneous expression of transdiagnostic constructs like anhedonia may reflect context-dependent adaptations to distinct underlying etiologies.

## Introduction

How do early experiences shape the way individuals navigate decisions that pit reward against threat? Early life adversity is a robust predictor of lifelong mental health burden (Kessler et al., 2010), contributing to depression (McGinnis et al. 2022) and suicidality (Dube et al. 2001). A growing body of work identifies unpredictability – the statistical structure of sensory and caregiving signals during sensitive developmental periods – as a particularly potent dimension of adversity (Baram et al. 2012; Glynn et al. 2019; Luby 2020; Xu et al. 2023; Birnie and Baram 2025). Unpredictability shapes neurocircuit maturation in ways that alter emotional processing and learning affected (Birnie & Baram, 2022). and neurobiological and computational evidence increasingly links these changes to specific disruptions in reward processing and uncertainty estimation (Davis & Glynn, 2024).

A recent efficient coding framework proposes that developing in unpredictable environments calibrates associative learning in ways that amplify subjective uncertainty, increase the weight of negative outcomes, and diminish the influence of positive outcomes (Harhen & Bornstein, 2024). These computational signatures overlap substantially with the phenomenology of anhedonia – a transdiagnostic symptom characterized by diminished capacity to experience or anticipate pleasure, in which impaired reward memory and an overall negativity bias are commonly observed (Dillon & Pizzagalli, 2018). Importantly, rodent models demonstrate that rearing under unpredictable conditions elicits anhedonic phenotypes, particularly in males (Birnie et al., 2023; Levis et al., 2021) and human research consistently links early-life unpredictability to anhedonia in adolescence and adulthood (Glynn et al., 2018; Hunt et al., 2024; Liu et al., 2025; Spadoni et al., 2022). Yet whether unpredictability and anhedonia exert the same or distinct influences on behavior, and whether their co-occurrence reveals or conceals underlying computational differences, has not been tested.

Decision-making in anhedonia has been studied primarily in tasks that isolate reward pursuit from punishment avoidance, yielding mixed results (Pizzagalli et al. 2005; Pizzagalli et al. 2008). However, the amplified uncertainty associated with early-life unpredictability and the negativity bias characteristic of anhedonia may be most consequential in situations of motivational conflict, where individuals must integrate competing reward-seeking and threat-avoidance drives under uncertainty. Approach-avoidance conflict tasks have been applied across several psychopathologies, including anxiety, depression, addiction, and PTSD (c.f. Letkiewicz et al., 2023) but findings have been inconsistent: both anxiety and depression are associated with greater decision uncertainty during approach-avoidance conflict (Smith et al., 2021), yet other work finds a specific avoidance bias in major depressive disorder (Pedersen et al., 2021). Critically, these inconsistencies may reflect unmodeled heterogeneity in the etiological backgrounds of participants, or unaccounted-for influences of trial-by-trial adaptation to task contingencies. This is precisely the kind of hidden structure that jointly modeling unpredictability and anhedonia could reveal.

Sequential decision problems offer a richer window into motivational conflict because they require individuals to weigh their current state, future prospects, and competing drives across multiple time steps, thus engaging a more naturalistic form of conflict resolution than elicited by single-shot full-information choice problems that directly reduce to more straightforward risk calculations (Korn & Bach, 2018; McNally, 2021). In a task developed by Korn & Bach (2018, 2019), participants navigated a multi-step foraging environment in which an optimal policy maximized the probability of survival across a finite time horizon by dynamically weighing present and distal reward or threat against the backdrop of continually-varying demands for survival. Participants differed in their adherence to this policy, with many trading off optimal choices for computationally simpler heuristics. This task is well suited to dissecting the influences of unpredictability and anhedonia because the optimal policy and specific deviations from it (e.g., a general bias toward or against approach, or excess threat sensitivity) can be separately quantified and linked to individual differences.

In this study, we adapt the Korn & Bach sequential decision-making task in two key ways to evaluate the influences of unpredictability and anhedonia on motivational conflict decision-making. We collect behavioral task data and validated survey instruments measuring self-reports of anhedonia and early life unpredictability from an online cohort. We first gamified the task in order to increase participant engagement and minimize individual differences in understanding task contingencies. Then, we optimized the task structure to allow identifying whether participants adhere to the optimal policy or deviate according to specific heuristics reflecting differing emphasis on competing motivational drives: either by a general bias for or against approaching reward, or a specific sensitivity to the likelihood of a threatening outcome over and above that prescribed by the optimal policy. Additionally, we exploit the structure of the task to identify maladaptive learning from unexpected negative outcomes. The new task, named *Revenge of the Zorn*, is described in **Figure 1**.

**Figure 1.**
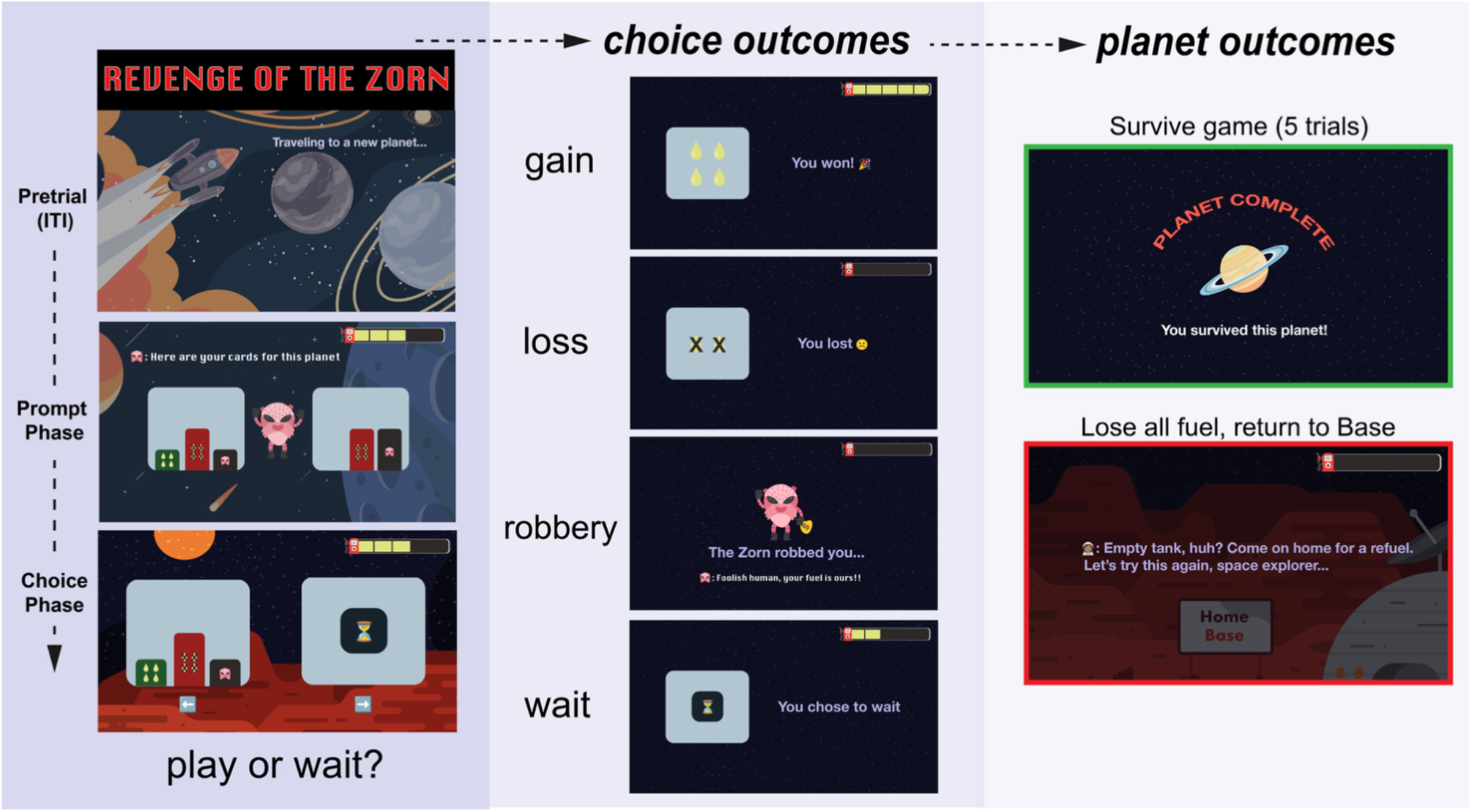
Revenge of the Zorn task gamification. Participants explore “planets,” where they are presented with opportunities to choose to “play” their card or decline to play the card, “wait.” Participants are presented with two cards of differing outcome probabilities, shown at random across 5 trials. Participants must play some number of the games to win fuel for their spaceship. Choice outcomes are gain, loss, wait (lose 1 unit) or robbed (lose all units). Participants can either survive a planet by completing all five of the card games (“days”), or lose all fuel and return to homebase, losing the planet.

We find that unpredictability and anhedonia exert opposing influences on choice: unpredictability *reduces* adherence to the optimal policy in a sex-dependent manner, while anhedonia *increases* it, partly through a bias toward cautious choices and heightened sensitivity to unexpected threatening outcomes. Mediation analysis reveals that anhedonia partially buffers the decision-making costs of early-life unpredictability. These findings suggest that these co-occurring conditions with shared etiological roots can nonetheless pull behavior in opposite directions. They additionally demonstrate that the heterogeneous behavioral expression of transdiagnostic constructs like anhedonia may reflect context-dependent responses to underlying adversity rather than uniform impairment.

## Results

### Participants generally play to the optimal policy, avoiding threat and slowing their response times as motivational conflict increases

Our primary hypothesis was that early-life unpredictability (ELU) and anhedonia effects would manifest as deviations from the optimal policy in favor of general (e.g. an overall tendency to forage or wait, independent of choice) or specific (e.g. an increased likelihood to wait at higher levels of threat, over and above that prescribed by the optimal policy) choice biases. We first examined the frequency with which participants chose to play their cards under increasing probability of survival according to the optimal policy (*p*_*forage*_**Fig 2a**). There was a significant relationship between optimal *p*_*forage*_ and the likelihood of playing the card (Kruskal-Wallis; χ^2^(2) = 738, p< .001); post-hoc tests indicated that each successive probability level was associated with greater proportion of play choices (Wilcoxon rank-sum: : low vs. medium, 12.4 ± 1.11, p < .001; medium vs. high, 34.5 ± 1.06, p < .001; low vs. high, 81.9 ± .581, p < .001). Similarly, we analyzed the effect of p(threat) alone, finding that the proportion of card-play trials decreased as the probability of threat (robbery by Zorn alien) increased (**Fig 2b**, (χ^2^(2) = 547, p< .001; low vs. medium, 54.5 ± .801, p < .001; medium vs. high, 39.1 ± .877, p < .001; low vs. high, 23.9 ± .852, p < .001).

**Figure 2.**
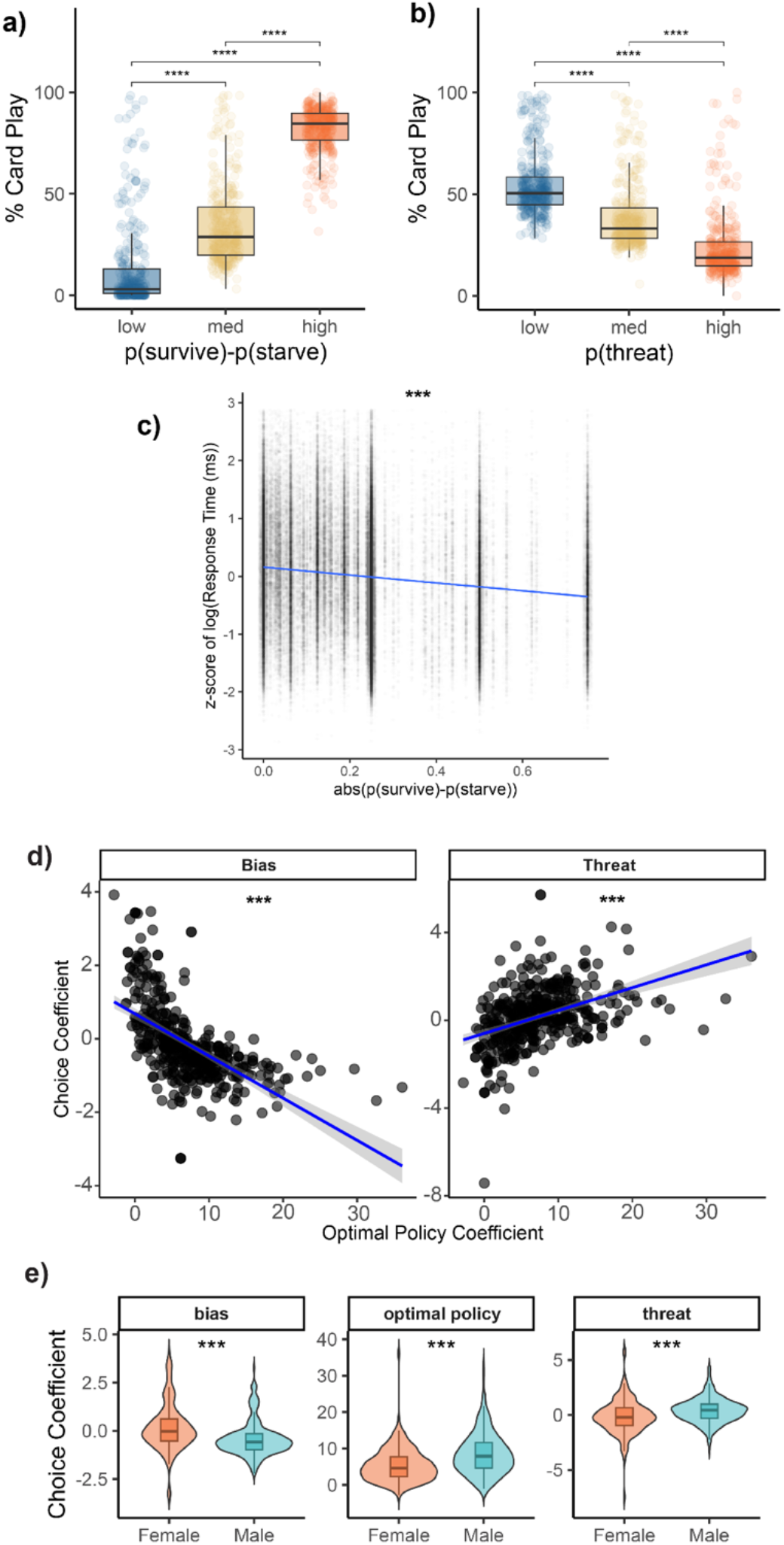
Overall gameplay analysis. A) Participants play their card with greater frequency when doing so results in a greater probability of surviving is greater than the probability of starving, as prescribed by the optimal policy (χ^2^(2) = 738, p<.001) B) Participants chose wait with greater frequency when the probability of threat was greater (χ^2^(2) = 547, p<.001). C) Response times were slowed when the prescription of play or wait by the optimal policy approached zero (F(1, 78174) = 1610, p <.001, SE bars included). D) The influence of optimal policy on choice is negatively associated with a bias towards playing (ρ = -0.68, S = 1 × 10^7^, n = 357, p<.001), while association to the effect of threat on choice is positive (ρ = 0.44, S = 4 × 10^6^, n = 357, p<.001). E) Females, on average, have a greater bias towards playing their card, less use of the optimal policy, and less card play (more waiting) under threat of robbery by Zorn predator.

To test whether participants deliberated longer when motivational conflict increased, we performed a linear regression analysis predicting log-transformed, z-scored response times from the absolute value of *p*_*survive*_ − *p*_*starve*_--the chance of survival or starvation resulting from the choice to “play” the card on the current day, assuming future days follow the optimal policy--a continuous measure of motivational conflict (**Fig 2c**), and including random effects to account for multiple observations per participant. The increase in the strength of the prediction of the optimal policy at each trial significantly decreased trial response times (β = -0.61, t(77,850) = -43, p < .001), indicating that participants responded more quickly when the motivational conflict decision was less ambiguous. In general, the mean optimal policy prescription in our task across all played trials favored wait (average *p*_*forage*_ = -0.091 ± .001, n trials = 93,700).

The individual threat β-coefficient and bias (intercept) parameter derived from the choice logistic regression described above (Methods) were correlated with the individual optimal policy β-coefficient to evaluate if the influences of these variables on choice were similar (**Fig 2d)**. A Spearman’s rank correlation was applied to test for correlations between choice coefficient variables with nonlinear relationships. This resulted in a significant, negative correlation between the optimal policy coefficient and the bias coefficient, where playing according to the optimal policy was associated with more waiting behavior (**Fig 2d left panel**, ρ = -0.68, S = 1 × 10^7^, n = 357, p<.001), consistent with widespread observations that individuals in sequential foraging settings “overharvest” – consistently forage more often than is optimal (Charnov 1976; Harhen and Bornstein 2023). Greater threat insensitivity (threat coefficient) was positively correlated with playing according to the optimal policy (**Fig 2d right panel**, ρ = 0.44, S = 4 × 10^6^, n = 357, p<.001). Sex differences in choice coefficients were found for the optimal policy choice coefficient, threat choice coefficient, and bias parameter **(Fig 2e)**. On average, males adhered more closely to the optimal policy (male: mean = 8.78 ± .407; females: mean = 5.70 ± .369; two-sample t-test: t(286) = 5.00, p <.001), were more likely to play their card under threat (male: mean = 0.451 ± .081; females: mean = -0.143 ± .119; two-sample t-test: t(285) = -4.00, p <.001), and had a baseline bias towards waiting overall, compared to females (male: mean = -0.458 ± .092; females: mean = 0.148± .063 two-sample t-test: t(286) = 5.00, p <.001).

### Early life unpredictability and anhedonia have opposing influences on optimal task choices

The effect of early life unpredictability on adherence to the optimal policy in task play was tested first in a linear model including all six of the derived latent factors from the QUIC survey and the optimal policy choice coefficient as the outcome. Only the General Unpredictability (F1) and Structure (F2) factors had a significant effect on participant adherence to the optimal policy **(Supp Table 2)**. Subsequent analyses focused on these two factors from the QUIC survey, including two separate linear regressions for each QUIC factor, plus sex and age as covariates and optimal policy choice coefficient as the outcome (**Fig 3a)**. Optimal policy use was significantly predicted by General Unpredictability when a sex interaction was modeled, where males had a significant drop in optimal policy use as the General Unpredictability measure increased (General Unpredictability:Males, β = -0.31, t(350) = -2.4, p = .015). Optimal policy use increased significantly across both sexes with Structure (β = 0.21, t(350) = 2.42, p = .016). Applying this model format to the sum of the QUIC survey (QUIC total) did not result in a significant association with adherence to the optimal policy (β = -0.037, t(350) = -0.99, p = .32). Taken together, this analysis is consistent with the recent theoretical proposal that early-life unpredictability increases sensitivity to uncertainty (Harhen & Bornstein, 2024), which we have previously shown to result in a decreased planning horizon in a reward-only foraging task (Chen et al. 2024), and which here manifests as a greater tendency to play the immediately presented card.

**Figure 3.**
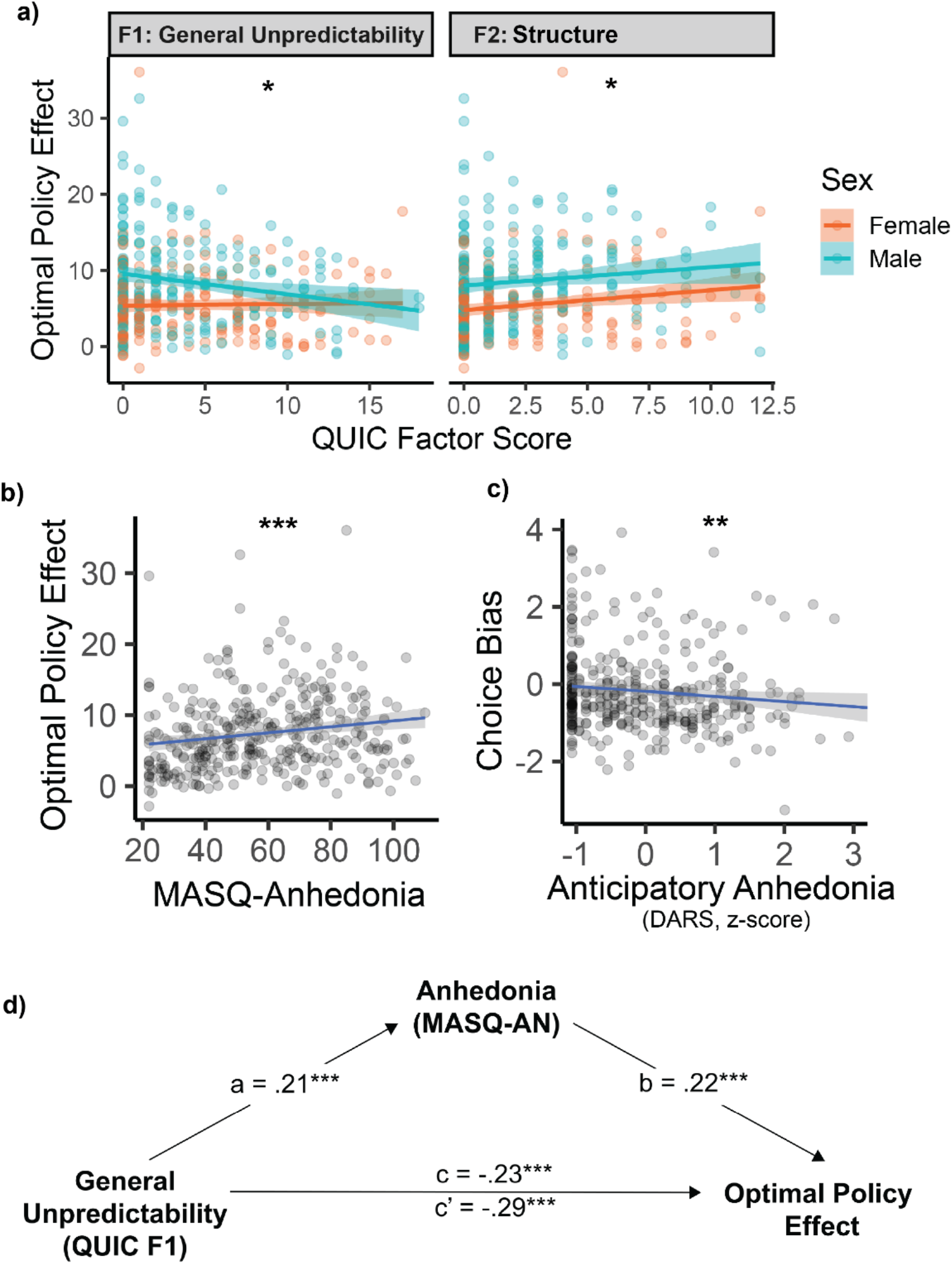
Early life adversity and anhedonia have opposing influences on optimal task choice. A) Unpredictability has a sex-dependent effect on optimal policy use, while structure has an opposing, but not sex-dependent, effect (panel 1. General Unpredictability:Males, β = -0.31, t(350) = -2.44, p = .015 ; panel 2: Structure, β = 0.21, t(350) = 2.42, p = 0.016). B) Anhedonia is associated with greater play according to the optimal policy (β = .467, t(351) = 3.80, p < .001). C) Anticipatory anhedonia is associated with more waiting overall for participants reporting greater anhedonia (DARS (inverted): β = -.150, t(351) = -2.82, p = .0051). D) Anhedonia partially mediated the relationship between General Unpredictability and optimal play (p<.001).

Next, the effect of anhedonia was also examined in a linear regression model, where the optimal policy choice coefficient was modeled as the outcome, and anhedonia summary measures from the MASQ, DARS, and SHAPS surveys were each modeled in separate regressions as predictors, along with sex and age effects (**Fig 3b)**. Anhedonic depression measured by MASQ-AD and SHAPS was significantly associated with greater adherence to the optimal policy following Bonferroni correction for multiple comparisons (MASQ: β = .467, t(351) = 3.80, p < .001; SHAPS (inverted): β = .118, t(351) = 3.02, p = .008), though the association with DARS did not survive multiple comparison correction (DARS (inverted): β = .56, t(351) = 2, p = .138). In a comparable analysis, the bias parameter was the outcome for a triplet of linear regression models with sex, age, and the three anhedonia measures described above. Anticipatory anhedonia, as measured by DARS, predicted a greater bias towards waiting choices at baseline, however, the other two measures did not survive multiple comparison correction (**Fig. 3c**, DARS (inverted): β = -.150, t(351) = -2.82, p = .0153, SHAPS (inverted): β = -.017, t(351) = -2.38, p = .162, MASQ-AD: β = -.00477, t(351) = -2, p = .138). Taken together, these findings are consistent with the idea that anhedonia works against the general tendency to overharvest, thus increasing adherence to the optimal policy, consistent with previous findings that major depression is associated with reduced patch leaving thresholds (Bustamante et al. 2024).

To evaluate further the opposing influences of unpredictability and anhedonia on optimal policy play in our task, we conducted a mediation analysis to assess if anhedonia (MASQ-AN) mediated the relationship between our latent construct of unpredictability (QUIC F1: General Unpredictability) and optimal play with using the mediation library in R and 5000 bootstrap samples **(Fig 3d)**. General Unpredictability significantly predicted less optimal play (β = -.18, t(355) = -3.54, p < .001). General Unpredictability significantly predicted more anhedonia (Path a: β = .21, t(355) = 4.09, p < .001). Anhedonia, the mediator, was associated with significantly more optimal play when controlling for unpredictability (Path b: β = .22, t(355) = 4.26, p < .001). In summary, higher unpredictability was associated with greater anhedonia, which in turn predicted increased engagement in the optimal survival strategy. This mediation analysis revealed a significant positive indirect effect of unpredictability on optimal play through anhedonia (.06, 95% CI: [.03, .1]) accounting for 25.5% of the total effect of unpredictability on survival strategy play. This analysis supports a partial mediation by anhedonia, as the direct effect of unpredictability on optimal play was still significant (Path c’: β = -.29, p < .001). A second moderated mediation analysis was performed to probe sex effects and found that while the direct effect of unpredictability on survival strategy play was moderated by sex, where males had a stronger association (Path c’ sex moderation: β = -.24, p < .001, the indirect (mediated) pathway through anhedonia was not **(Supp Fig 3;** Path a sex moderation: β = .11, t(351) = 1.30, p = .195); Path b sex moderation: β = .07, t(351) = .53, p = .59).

### Anhedonia may promote adaptive behavior following unexpected threat outcomes

To investigate the effect of unexpected predation events on subsequent choices, a threat prediction error (TPE) value was computed and input as a predictor to a choice logistic regression, as described above (Methods). Kruskal-Wallis testing of the choice coefficients for the three subsequent planets indicated a significant effect of TPE on choice (**Fig 4a**. χ^2^(2) = 105, p< .001). Post-hoc Wilcoxon rank sum testing with Bonferroni correction showed that TPE increased the tendency to choose “wait” strongly on the first planet following robbery, as well as on the third planet following the robbery event (n+1 mean: -.367 ± .048, z = -11.7, p < .001; n+2 mean: -0.005 ± 0.022, z = -0.75, p = .5; n+3 mean: -.067 ± 0.035, z = -3 p = .003).

**Figure 4.**
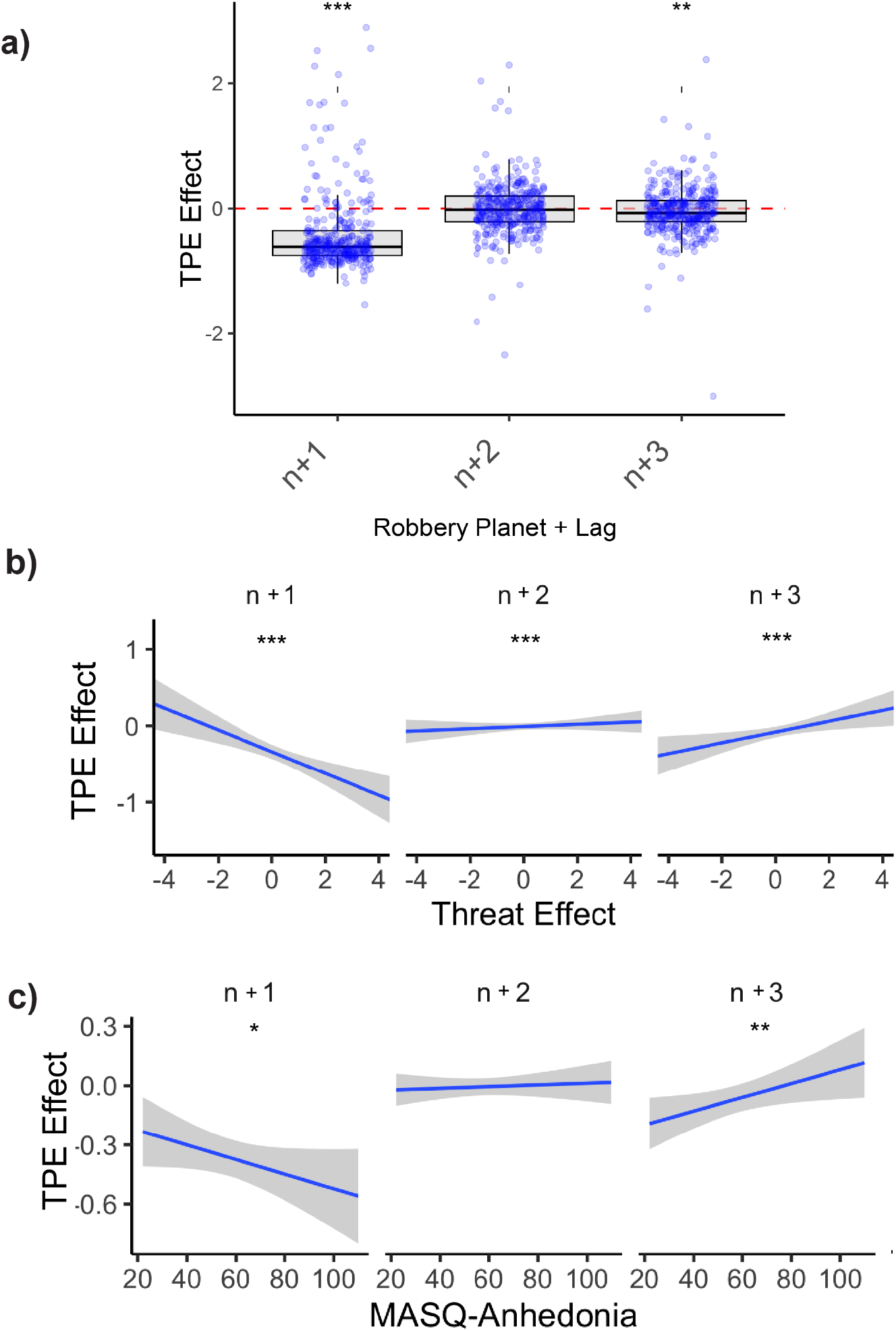
Effects of unexpected robbery outcome on following choices. a) Following an unexpected robbery, participants were more likely to wait in their average responses, with a strong effect on responses during the planet immediately following the robbery event (n+1 mean: -.367 ± .048, z = -11.7, p < .001; n+2 mean: -0.005 ± 0.022, z = -0.75, p = .5; n+3 mean: -.067 ± 0.035, z = -3 p = .003). B) Participants who were more likely to play under threat had the greatest increase in waiting following an unexpected robbery on the following planet (n+1, β = -.14, t(1065) = -5.2, p < .001), but this effect reversed in the second and third planets (n+2*threat effect,, β = .15, t(1065) = 4.7, p < .001, n+3*threat effect, β = .21, t(1065) = 5.5, p < .001). C) Participants with greater levels of anhedonia were more likely to wait on the first planet trials immediately following the unexpected robbery (n+1, β = -.003, t(1065) = -2.2, p =.02), but this effect also decreased significantly by the second (n+2*MASQ-AD, β = .004, t(1065) = 1.79, p =.07) and reversed direction by the third planet (n+3*MASQ-AD, β = .007, t(1065) = -5.2, p =.002).

The TPE choice coefficients were then entered as outcomes into separate linear regression models where the planet-lag was placed in an interaction with each predictor of interest (**Fig 4b & c)**. The first linear regression model included the threat choice coefficient as a predictor, analyzing if one’s sensitivity to threat might predict behavior following an unexpected robbery (as measured by the TPE coefficient). Participants who were more likely to play their card under threat were the most likely to wait following an unexpected robbery on the first following planet (**Fig 4b**. n+1, β = -.14, t(1065) = -5.2, p < .001), but at the second and third planet, the relationship reversed, where participants were again more likely to play their card (n+2*threat effect, β = .15, t(1065) = 4.7, p < .001, n+3*threat effect, β = .21, t(1065) = 5.5, p < .001). We note that the n+1 effect may partly reflect regression toward the mean, as participants with stronger baseline threat approach have more room to shift toward avoidance following a surprising negative outcome; the subsequent reversal at n+2 and n+3 is consistent with this interpretation but could also reflect genuine recovery of baseline threat tolerance.

MASQ-AD was entered in a new linear regression model (**Fig 4c)**, and shared a similar relationship as threat effect, where anhedonia was associated with greater waiting on the subsequent planet (n+1, β = -.003, t(1065) = -2.2, p =.02), but that effect weakened and reversed on the second (n+2*MASQ-AD, β = .004, t(1065) = 1.79, p =.07) and third following planets (n+3*MASQ-AD, β = .007, t(1065) = -5.2, p =.002). To test the relationship between early life unpredictability and the effect of an unexpected threat outcome on choice, we tested General Unpredictability (QUIC F1) in a third linear regression model, where participants with greater childhood unpredictability were marginally less likely to wait on the first planet following an unexpected robbery event (n+1, β = .016, t(1065) = 1.93, p =.05), and this effect reversed on second planet, though marginal (n+2*General Unpredictability: β = -.021, t(1065) = -1.66, p =.098) and no effect was observed on the third planet (n+3; β = -.015, t(1065) = -1.23, p =.21). A follow-up sex-interaction analysis for the n+1 planets found that males with unpredictability were less likely to wait following unexpected robbery (**Fig 5**, General Unpredictability*Sex(Males): β = .066, t(351) = 2.94, p =.003).

**Figure 5.**
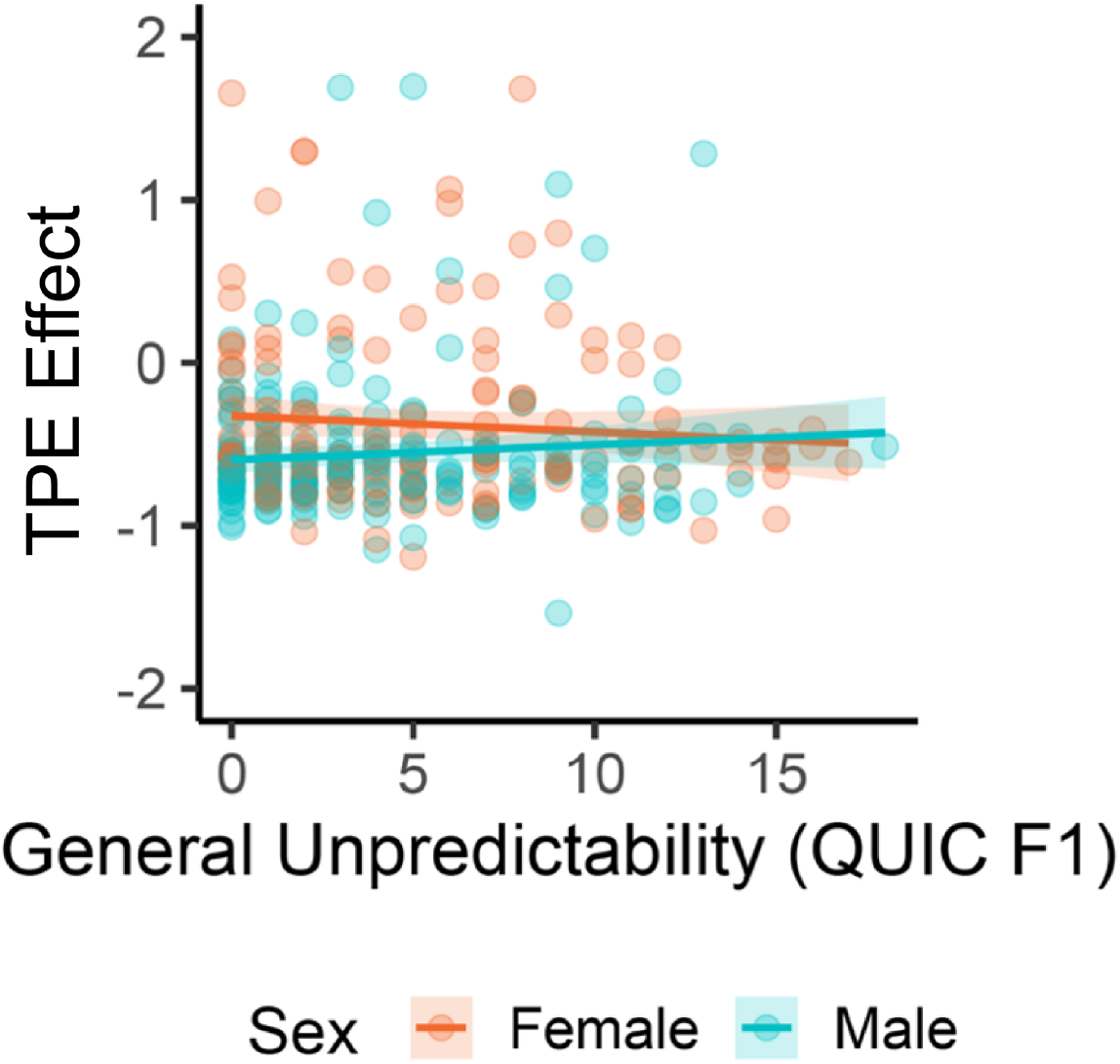
Sex-dependent effect of unpredictability on behavior following unexpected threat outcome. Males with higher levels of unpredictability were less likely to choose “wait” on the trials immediately following an unexpected robbery (General Unpredictability*Sex(Males): β = .066, t(351) = 2.94, p =.003).

### Participants’ sensitivity to threat changes across the task in temporal analysis

To evaluate fluctuations in participant threat sensitivity across the task, the threat choice coefficient was compared across the three task blocks (20 planets each, **Supp Fig 4**). The average effect of threat on choice differed significantly across the three blocks (**Supp Fig 4a**, Kruskal-Wallis test: χ^2^(2) = 105, p< .001). Post-hoc pairwise Wilcoxon tests revealed that threat sensitivity was lowest in the middle block, where participants were most likely to play their card under threat (.61 ± .09), but the middle block did not differ significantly from the first block (.13 ± .09, p < .001). The effect reversed in direction in the late block, where they were most threat sensitive and most likely to choose “wait” under threat, and this differed significantly from both the early and middle planet blocks (-.28 ± .08, late vs. early: p = .02; late vs. middle: p <.001).

We next evaluated the effect of early life unpredictability and anhedonia on threat choice coefficients across each block of the task. The reduced number of trials lead to convergence issues for some participants. As a result, two participants were completely removed as outliers, and 58 participants removed as partial outliers (middle and late blocks had failure of convergence; respective individual coefficients for those blocks were removed from analysis). All outliers in this analysis were removed using the Tukey method (outside 1.5 times the interquartile range). General Unpredictability (QUIC F1) significantly predicted card play under threat in the early block (**Fig 4b**, β = .027, t(332) = 2.27, p =.024), but not the middle (β = .005, t(332) = .49, p =.62) nor late blocks (β = - .006, t(332) = -.59, p =.56). Anhedonia (MASQ-AD) had only marginal effect on threat choice coefficient in the first block, and no effect in subsequent blocks (**Fig 4c**, early: β = .007, t(332) = 1.69, p =.09, middle: β = -.005, t(332) = 1.40, p =.16, late: β = -.005, t(332) = -1.42, p =.17). Threat sensitivity increased as the task progressed under early life unpredictability, but not in anhedonia.

## Discussion

In this study, we examined the performance of individuals in a motivational conflict challenge embedded in a sequential decision-making task under threat of predation. We found that participants generally played the task optimally, behaving in a manner consistent with optimizing their survival probability across five simulated “days” on each planet. This tendency was modulated in individuals with experiences of elevated early-life unpredictability (unpredictability), and those who exhibited strong symptoms of anhedonia according to validated self-report instruments. Interestingly, these experiences and attitudes, which are correlated with each other in our sample and more generally in the population, had opposing effects on adherence to the optimal policy. Anhedonia supported more optimal play under motivational conflict, while unpredictability had a sex-dependent effect on optimal play, such that males showed less adherence to the optimal policy. One manner in which anhedonia improved adherence to the policy was through its association with an overall choice bias towards waiting. Additionally, participants who reported greater anhedonia symptoms also showed greater sensitivity to surprising predation experiences, as did those who were less sensitive to threat on average. Childhood unpredictability portended greater threat insensitivity early in the task, though this moderated with further play.

The sex differences we observed were consistent with previous findings: males adhered more to the optimal policy on average, possibly driven by a greater choice bias towards waiting overall. However, males were more likely to play their card under elevated threat than were females, consistent with previous research finding fewer approach choices in females in an AAC task under threat of a negative affective stimulus (Aupperle et al., 2011).

Early life structure increased adherence to the optimal policy overall, while males who reported higher levels of childhood unpredictability were less adherent to the optimal policy. Unpredictability in childhood has been correlated with poorer cognitive and mental health outcomes in childhood and adolescence, which may persist until adulthood (Glynn et al., 2018; McGinnis et al., 2022). Furthermore, a sex-specific effect of unpredictability is supported by rodent model literature, where early life experiences are differentially encoded neurally by sex (Kooiker et al., 2023), as well as human findings of differences in BOLD responses to novelty between sexes, with males expressing diminished limbic BOLD activation (Davis et al., 2025). A sex-dependent difference in the neurodevelopmental effects of unpredictability may underlie a greater impairment of optimal play for males reporting childhood unpredictability.

Anhedonia, by contrast, was associated with greater adherence to the optimal policy and a stronger bias toward “wait” choices. Previous research has linked decreased approach behavior to major depressive disorder (Pedersen et al., 2021), a pattern typically interpreted as maladaptive. However, in the context of our task engaging putative evolutionarily conserved sequential choice behavior, this same tendency corrects the widely observed bias towards overharvesting (“play” choices) in foraging settings (Blanchard and Hayden 2015), yielding decisions more closely aligned with the survival-optimal policy. This does not imply that anhedonia is globally adaptive. Its debilitating effects on quality of life across psychiatric conditions are well established. Rather, our findings suggest that the behavioral recalibrations associated with anhedonia may be context-dependently beneficial: in environments where reward pursuit carries genuine risk, a dampened approach drive and heightened threat sensitivity can improve decision quality. This interpretation aligns with computational accounts of consummatory anhedonia in which “belief warping” (modifications to an individual’s internal state that attenuate hedonic valuation) serves as a response to chronic stress (Hall et al., 2024), and with the recent proposal that such adjustments may serve a compensatory function in the presence of the amplified uncertainty and negativity bias produced by early-life unpredictability (Zhang and Bornstein 2026).

As previously documented (Glynn et al., 2018; Hunt et al., 2024; Liu et al., 2025; Spadoni et al., 2022), early life unpredictability was associated with greater anhedonia symptoms in our sample. A mediation analysis revealed that anhedonia partially accounted for the relationship between unpredictability and adherence to the optimal policy. This pattern is consistent with a model in which anhedonic attitudes buffer the decision-making costs of early-life unpredictability. However, important caveats apply here. Our data are cross-sectional, and mediation analysis in this context cannot establish the temporal or causal ordering of variables; the observed statistical mediation is also consistent with alternative causal structures, including shared underlying factors that independently influence both anhedonia and decision behavior. Longitudinal or experimental designs will be needed to determine whether anhedonia causally mediates the effects of early-life unpredictability on decision-making, or whether both reflect parallel downstream consequences of altered neurodevelopment.

One pathway through which anhedonia may be associated with more optimal play in this task is through greater negative feedback sensitivity, which could allow for trial-by-trial adjustments that counteract initial choice biases. This possibility is consistent with evidence that anhedonia is accompanied by an outsized influence of negative prediction errors on mood, potentially reflecting a negative attentional bias driven by amygdala sensitization that outweighs a less responsive dopaminergic reward system (Dillon & Pizzagalli, 2018; Harhen & Bornstein, 2024; Lim et al., 2012). In our data, sensitivity to threat was similarly associated with more “wait” choices on the first planet following a surprising predation event. Sex differences were also apparent in this domain: males with greater childhood unpredictability were less likely to wait following unexpected threat outcomes. If this pattern reflects a sex-specific insensitivity to negative feedback, it could contribute to the reduced optimal play observed in this group. However, establishing this link will require designs that can isolate the temporal dynamics of feedback-driven choice adaptation.

The influence of threat probability differed across blocks of the task, and childhood unpredictability resulted in significant threat tolerance in the early block, which is related to less optimal play. Unpredictability is associated with greater risky and externalizing behaviors (Simpson et al. 2012; Doom et al. 2016; Srivastava et al. 2025), though it is worth noting that the threat tolerance found in our task disappears by the second tercile of trials. Few studies have examined the effects of early-life unpredictability on learning rates, but this result may suggest that high-unpredictability participants are slower to apply the optimal strategy due to slower threat learning, also consistent with a predicted increase in subjective uncertainty (Harhen & Bornstein, 2024). In contrast to unpredictability, previous work has examined differences in learning where individuals with high scores on anhedonia surveys differentially adapt their choice strategies across reward-choice tasks (Pizzagalli et al., 2005, 2008). Together, these findings highlight the key role of between-trial adaptation dynamics in understanding the impacts of anhedonia and early-life unpredictability on threat sensitivity, even in situations where full information limits opportunities for learning.

In summary, early-life unpredictability and anhedonia exert opposing influences on motivational conflict decision-making: unpredictability shifts behavior away from the optimal policy in a sex-dependent manner, while anhedonia promotes adherence to it through dampened approach and heightened threat sensitivity. That these co-occurring conditions pull behavior in opposite directions has implications both for how we study their effects jointly rather than in isolation, and for how we interpret the heterogeneous expression of transdiagnostic symptoms more broadly.

Several limitations warrant consideration. Our design is cross-sectional and observational, precluding causal claims about the relationships among early-life unpredictability, anhedonia, and decision behavior. Anhedonia was assessed via validated self-report instruments rather than clinical evaluation, and participants were recruited online, limiting experimental control. Future work could extend the task in several directions: introducing ambiguous threat probabilities (Levy et al., 2010), unexpected rewards, or graded rather than sudden-death predation outcomes would test the generality of the present findings across different motivational conflict structures. Post-task strategy reports could clarify whether the behavioral signatures we observe reflect deliberate policies or emergent biases. Neuroimaging implementations could evaluate the contributions of regions implicated in motivational conflict, including the paraventricular nucleus of the thalamus (McNally, 2021) and hippocampus (Bach et al., 2014), and longitudinal designs tracking individuals from childhood could establish whether the mediating role of anhedonia reflects a developmental process.

The approach developed here, namely decomposing threat sensitivity, choice bias, and sequential adaptation patterns within a single task, could be applied to other conditions in which motivational conflict is implicated, including anxiety, PTSD, and attention deficits. These conditions frequently co-occur, and their presence together or in isolation may give rise to distinct computational profiles, as we demonstrated here for unpredictability and anhedonia. More broadly, our findings raise the possibility that the heterogeneous behavioral expression of transdiagnostic constructs reflects, in part, the interaction of a given symptom with the etiological context from which it emerged: anhedonia following early-life unpredictability may carry different behavioral signatures than anhedonia arising from other pathways. If so, interventions targeting anhedonic symptoms (Hall et al., 2024) may need to account for the possibility that reducing anhedonia could unmask distinct maladaptive patterns tied to the underlying etiology (Zhang and Bornstein 2026) rather than uniformly improving function.

## Methods

### Participants

Four-hundred participants were recruited across 35 countries from the Prolific database (www.prolific.com, 2025) and collected between the months April - October, 2025. The final sample consisted of 357 participants following quality assurance filtering, including failed attention checks during survey data collection and playing >80% of trials in task (**Table 1**, 157 F, 198 M, 2 “Other”; mean ± SD: 32 ± 13 years of age).

**Table 1.**
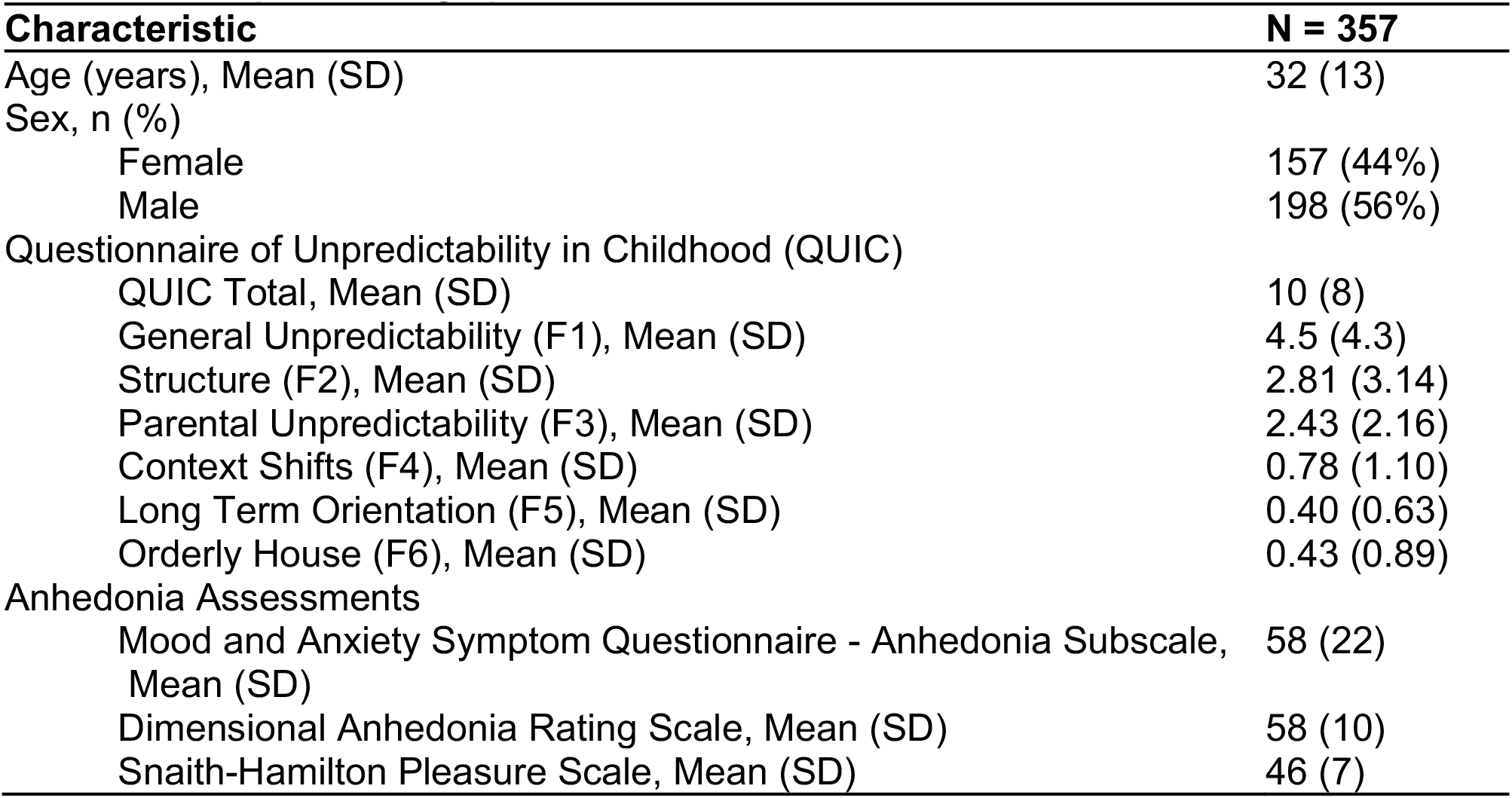
Participant Demographics.

| Characteristic | N = 357 |
| --- | --- |
| Age (years), Mean (SD) | 32 (13) |
| Sex, n (%) |  |
| Female | 157 (44%) |
| Male | 198 (56%) |
| Questionnaire of Unpredictability in Childhood (QUIC) |  |
| QUIC Total, Mean (SD) | 10 (8) |
| General Unpredictability (F1), Mean (SD) | 4.5 (4.3) |
| Structure (F2), Mean (SD) | 2.81 (3.14) |
| Parental Unpredictability (F3), Mean (SD) | 2.43 (2.16) |
| Context Shifts (F4), Mean (SD) | 0.78 (1.10) |
| Long Term Orientation (F5), Mean (SD) | 0.40 (0.63) |
| Orderly House (F6), Mean (SD) | 0.43 (0.89) |
| Anhedonia Assessments |  |
| Mood and Anxiety Symptom Questionnaire - Anhedonia Subscale, Mean (SD) | 58 (22) |
| Dimensional Anhedonia Rating Scale, Mean (SD) | 58 (10) |
| Snaith-Hamilton Pleasure Scale, Mean (SD) | 46 (7) |

### Task Design

Participants played “Revenge of the Zorn”, a gamified adaptation of a virtual motivational conflict task (Korn & Bach, 2019). Participants have the goal of exploring planets (60 planets in total) while minimizing the probability of running out of fuel (i.e., probability of starvation), for all five days (5 trials) on each planet and also avoiding sudden-death by Zorn alien robbery of fuel (i.e., threat of predation). At the start of each planet, participants were given two “cards” each depicting a probability of gaining a given number of fuel points, a probability losing a given number of fuel points, and a probability of robbery, which would result in an empty fuel tank and loss of planet (**Fig 1**. Prompt phase). The two cards given at the start of each planet carry forward across all trials of the current planet in a predetermined sequence. At the start of each trial, the participant was presented with one of the cards, and asked whether they decided to “play” in which case that card’s probabilities were used to generate the outcome, or to “wait,” in which case they remained safe but incurred the sure loss of one fuel point. Because the “wait” option costs one fuel point, participants were not able to survive by simply choosing to “wait” for all five days on the planet, and instead had to weigh the risks of ensuring survival under the current choice set against the possibility of alternative contingencies should the other card be drawn. Participants reviewed task instructions, which were read to them by a recorded voice, and were given a short quiz to review the most important information. Then, a practice round of five planets was played before proceeding to the main task, hosted on Pavlovia (www.pavlovia.org). Practice planets were selected to give individuals experience with planets that elicited both maximally easy/difficult choices (low/high conflict), as well as intermediate levels.

### Planet selection and optimal policy derivation

The authors of the original version of this motivational conflict task modeled numerous decision-making heuristics for behavioral analysis within this task (Korn & Bach, 2019). Here, our design goal was to evaluate choices when the reward-seeking drive (the *optimal policy*) and pure predation-avoidance (*hard-avoid*) were at odds. In MATLAB (Mathworks, Inc., R2024b), we first constructed planets (two-card pairs, presented at the start of each planet) optimized such that these two strategies would be in conflict at a range of values. Backwards induction was used to compute the probability of ultimately running out of fuel (“starving”; *p*_*starve*_) as a result of making each choice (i.e., wait or play), on each day, under uncertainty. The formal variables of interest for each trial are as follows: *c*_*f*_, the cost of foraging (e.g., the loss); *p*_*f*_, the probability of winning; *g*_*f*_, the amount to be gained; *p*_*r*_, the probability of being robbed; *c*_*w*_, the cost of waiting (fixed at -1). First, we calculated (*p*_*starve*_) for each option (Play, Wait) on Day 5, given two choice options (cards) on this planet ([*p*_*r*_, *g*_*f*_, *p*_*f*_, *c*_*f*_]) and possible fuel levels [1:5]. We assume that the agent believes there is a 50% probability of each card on each day, as participants were instructed, and that the agent prefers the option that minimizes *p*_*starve*_. We computed *p*_*forage*_, the probability that the agent would choose to play their card, as a logistic transform of the difference between *p*_*starve*_ values for each option:

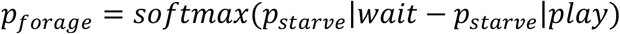

*p*_*wait*_ was computed as 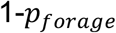, and the motivational conflict level of each trial was computed as the difference in probability of survival between the two choices, and later computed for each day (trial) on the planet. Inverse temperature was set to 1 for all simulations–we later validated, in our experiment sample, that the resulting probabilities are a good match for participant deliberation by comparing the indifference point inferred from response times to the one implied by this calculation (Supplemental Figure 1). Simulations of all planet combinations were computed for *p*_*r*_ = [.25, .5, .75], *g*_*f*_ = [1, 2, 3, 4, 5], *p*_*f*_ = [.25, .5, .75], *c*_*f*_ = [−1, −2, −3, −4]. Planets were then selected for a uniform distribution of *p*_*forage*_ values for use in the task such that no dominant strategy (reward-seeking vs. hard-avoid) had a better chance of survival and for a minimal correlation between the two strategy choices.

The final game sequence of planets and corresponding card choices were selected for use in the task, and the optimal policy *p*_*forage*_ values for each choice option on all card, planet, and fuel combinations were computed. We then matched each of the participants’ real trials to the corresponding *p*_*forage*_ value (“optimal policy” value), where larger values indicated a higher probability for survival if the card was played, and lower values correspond to a higher probability of survival if “wait” was chosen.

### Choice logistic regression

Game variables (optimal policy and threat probability) were evaluated for their effect on participants’ choices, first using a logistic regression to compute for each individual the effects of the optimal policy (*p*_*forage*_) and threat probability (*p*_*threat*_) on choice.

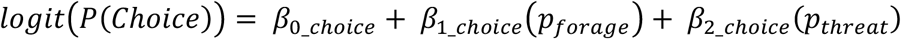

The participant-level coefficients resulting from this regression were later entered into linear regression models to investigate their relationship to response times, early life unpredictability, and anhedonia. Linear regression models were fitted in R (version 4.5.1), using the lme4 package, (Bates et al. 2015).

### Negative feedback sensitivity

Previous work has suggested that ELU leads to outsized influence of surprising negative outcomes on subsequent behavior (Harhen and Bornstein 2024), and that anhedonia is associated with altered adaptation of response tendencies across a task with randomized outcomes (Dillon and Pizzagalli 2018). Here, we examined how surprising negative outcomes alter choices across trials, even in the presence of full information at each trial (i.e. no explicit learning). A threat prediction error (TPE) was calculated to evaluate how negative feedback from an unexpected robbery event affected choices on the subsequent planets. The difference between the outcome (1/0 binary representing yes/no robbery) and the probability of the Zorn alien robbing (threat probability) was computed for each trial and used as a predictor in a choice probability logistic regression. The participant-level TPE estimates were later used as outcomes in linear regression models to compute the effect of early life unpredictability and anhedonia on learning from surprising negative outcomes.

### Assessment of Unpredictability and Anhedonia

Participants completed nine surveys assessing demographics, childhood environment, and anhedonia symptoms. Early life unpredictability was assessed using the 38-item Questionnaire of Childhood Unpredictability (QUIC) (Glynn et al., 2019). Anhedonia was assessed using the Mood and Anxiety Symptom Questionnaire Anhedonia subscale, which characterizes anhedonic depression (MASQ-AN, (Bredemeier et al., 2010; Watson & Clark, 2012)). The Dimensional Anhedonia Rating Scale (DARS, 17 items) and the Snaith-Hamilton Pleasure Scale (SHAPS, 14 items) were also completed by the participant to assess anticipatory and consummatory anhedonia symptoms (Rizvi et al., 2015; Snaith et al., 1995). To adjust scales so that greater scores were consistent with more anhedonia, DARS and SHAPS scores were inverted and scaled to a z-score for use in later analyses.

### Early life unpredictability exploratory factor analysis

Early-life unpredictability is a multi-dimensional construct, with some aspects endorsed by some individuals and not others. We examined whether latent factors of this construct are differentially related to the outcome measures of interest in our analysis. To accomplish this, we conducted an exploratory factor analysis to identify factors related to distinct aspects of childhood environmental unpredictability measured by the QUIC assessment, using JASP (version 0.19.3; JASP Team (2025)). Data collected in a separate sample (n = 735; collected on Amazon Mechanical Turk between 02/22/2021 and 04/04/2024 using the CloudResearch Approved Participants filter) was used to compute factors. Maximum likelihood method was employed, and an oblique rotation (Promax) was applied since data from questions of the QUIC were expected to correlate. The number of factors was determined using a threshold of eigenvalues > 1.0 (Kaiser criterion), and factor loadings greater than 0.4 were accepted. Six factors were identified (**Table 2)**. Sums for each factor score accounted for reverse-coded items.

**Table 2.**
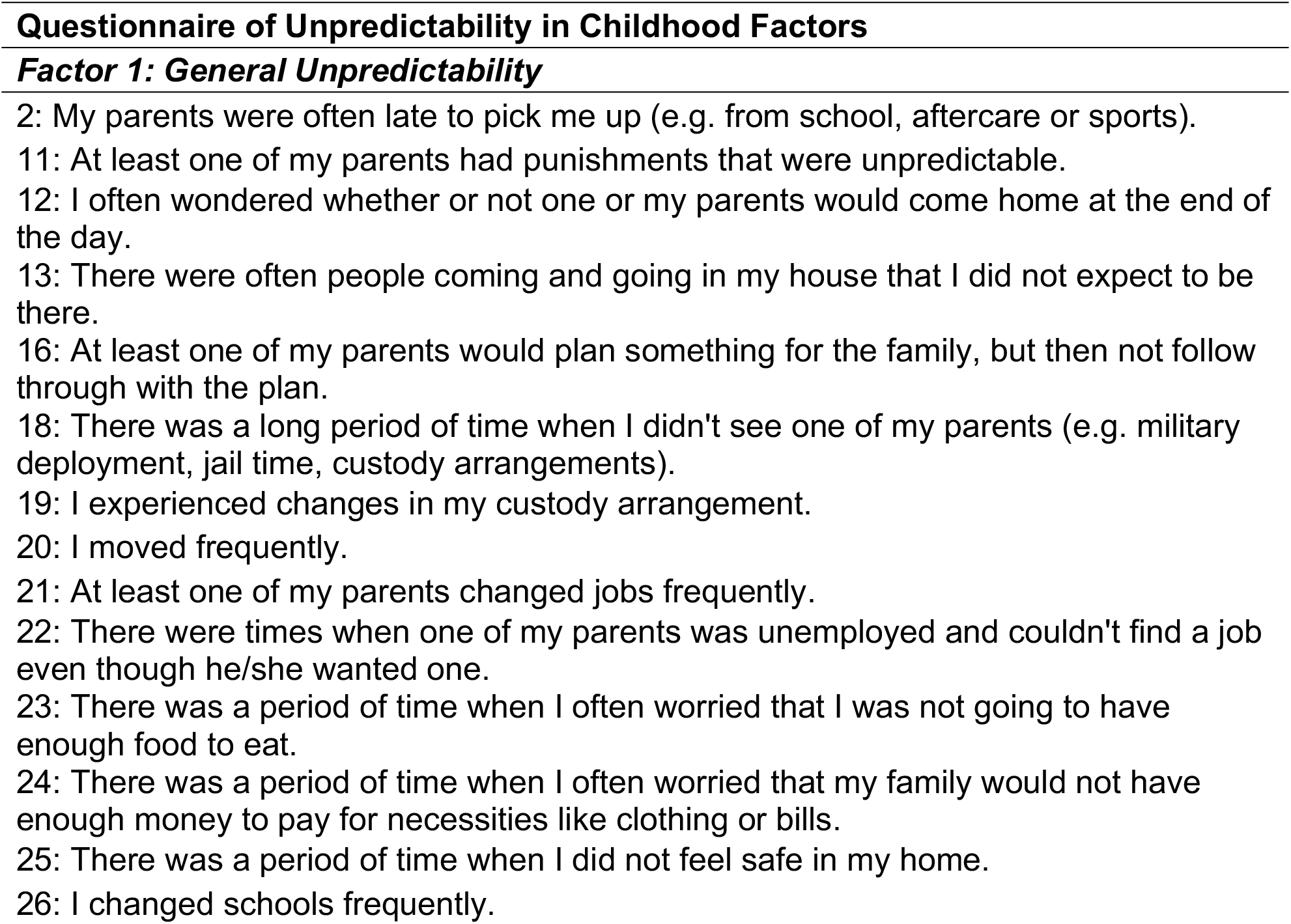
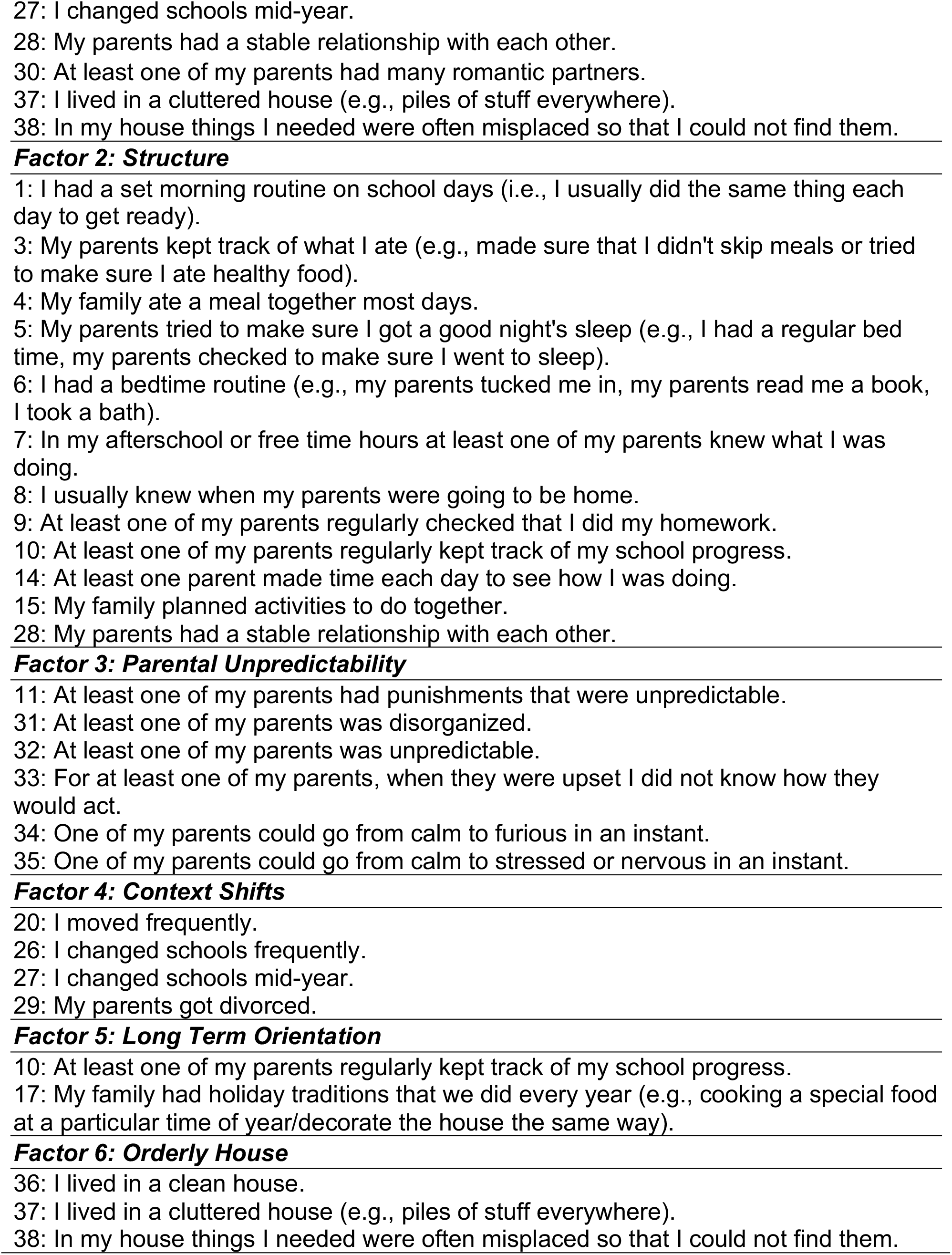
QUIC Factor Scores.

## Acknowledgements

The authors would like to acknowledge support from NIMH R01MH128306 (MAY, AMB), P50MH096889 (TZB, HSS, LMG, EPD, MAY), and T32MH119049 (BTL).

## Data and code availability

Upon publication all anonymized data, analysis scripts, and experiment code will be made freely available at the authors’ GitHub.

