## Supplementary material for "Anhedonia buffers the effects of early-life unpredictability on threat-reward decision-making"

### Extended Data

#### Supplementary Table 1: Bivariate correlations between study variables

| Variable | 1. | 2. | 3. | 4. | 5. | 6. | 7. | 8. | 9. | 10. | 11. | 12. | 13. |
|---|---|---|---|---|---|---|---|---|---|---|---|---|---|
| **1. Age** | — | | | | | | | | | | | | |
| **2. QUIC Total** | 0.07 | — | | | | | | | | | | | |
| **3. General Unpredictability (F1)** | 0.04 | 0.89*** | — | | | | | | | | | | |
| **4. Structure (F2)** | 0.08 | 0.78*** | 0.45*** | — | | | | | | | | | |
| **5. Parental Unpredictability (F3)** | 0.06 | 0.75*** | 0.63*** | 0.41*** | — | | | | | | | | |
| **6. Context Shifts (F4)** | 0.04 | 0.64*** | 0.69*** | 0.40*** | 0.33*** | — | | | | | | | |
| **7. Long Term Orientation (F5)** | 0.03 | 0.55*** | 0.29*** | 0.70*** | 0.30*** | 0.24*** | — | | | | | | |
| **8. Orderly House (F6)** | 0.05 | 0.56*** | 0.48*** | 0.42*** | 0.38*** | 0.24*** | 0.21*** | — | | | | | |
| **9. MASQ Anhedonia** | 0.06 | 0.37*** | 0.21*** | 0.40*** | 0.32*** | 0.16** | 0.30*** | 0.24*** | — | | | | |
| **10. DARS Total** | -0.05 | -0.15** | -0.08 | -0.20*** | -0.09 | -0.01 | -0.18*** | -0.16** | -0.32*** | — | | | |
| **11. SHAPS Total** | -0.04 | -0.31*** | -0.16** | -0.38*** | -0.22*** | -0.10 | -0.27*** | -0.22*** | -0.63*** | 0.51*** | — | | |
| **12. Survival** | -0.02 | -0.05 | -0.18*** | 0.11* | -0.05 | -0.08 | 0.06 | 0.02 | 0.17** | -0.09 | -0.16** | — | |
| **13. Threat** | -0.10* | -0.00 | -0.06 | 0.03 | 0.04 | -0.02 | 0.02 | -0.00 | 0.05 | -0.01 | -0.08 | 0.41*** | — |
| **14. Bias** | 0.06 | -0.00 | 0.06 | -0.08 | -0.00 | 0.06 | -0.04 | -0.02 | -0.08 | 0.13* | 0.12* | -0.58*** | -0.60*** |

N = 357, Pearson's *r*.
*p < .05, ** p < .01, *** p < .001

**Supplementary Table 2: Linear Regression Model: QUIC Factor Scores Effect on Optimal Policy Adherence**

| Variable | *B* | *SE* | *t* | *p* |
|---|---|---|---|---|
| ***Intercept*** | ***7.876*** | ***0.454*** | ***17.352*** | ***< .001**** |
| **General Unpredictability (F1)** | **-0.553** | **0.116** | **-4.757** | **< .001***** |
| **Structure (F2)** | **0.348** | **0.143** | **2.434** | **0.015*** |
| Parental Unpredictability (F3) | 0.203 | 0.173 | 1.170 | 0.243 |
| Context Shifts (F4) | 0.476 | 0.365 | 1.302 | 0.194 |
| Long Term Orientation (F5) | -0.163 | 0.635 | -0.257 | 0.798 |
| Orderly House (F6) | 0.614 | 0.383 | 1.604 | 0.110 |

*Note.* $N = 355$. Model fit: $R^2 = 0.092$, Adjusted $R^2 = 0.076$, $F(6, 348) = 5.85$, $p < .001$. † $p < .10$. * $p < .05$. ** $p < .01$. *** $p < .001$.

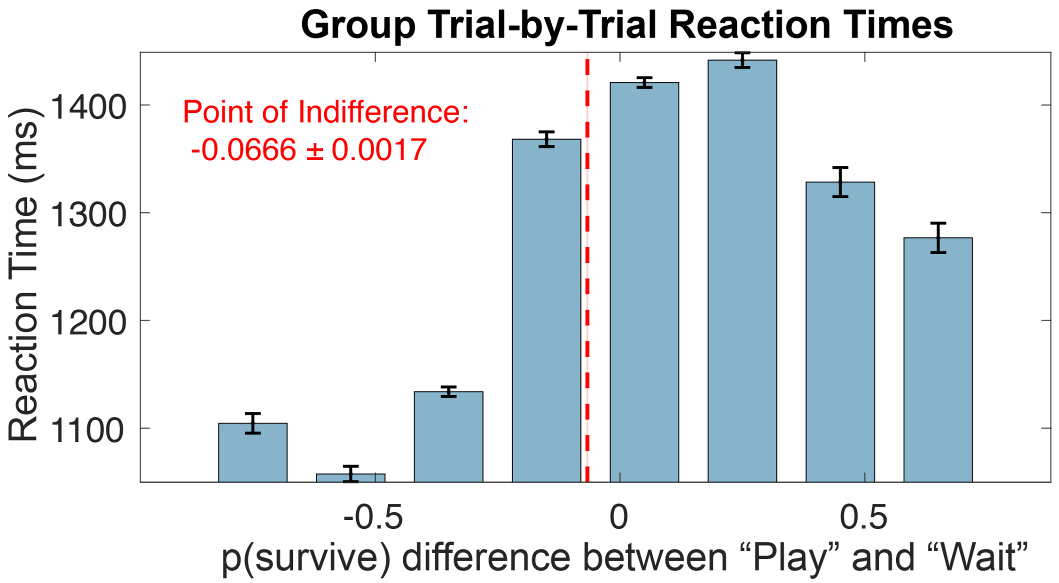


**Supplemental Figure 1. Motivational conflict values are reflected in participant reaction times.** Participants' average point of indifference approximates zero when difference in p(survive) between choices nears zero. Point of indifference is calculated by the sum of trial reaction times multiplied by trial difference in p(survive) between choices divided by the sum of trial reaction times.

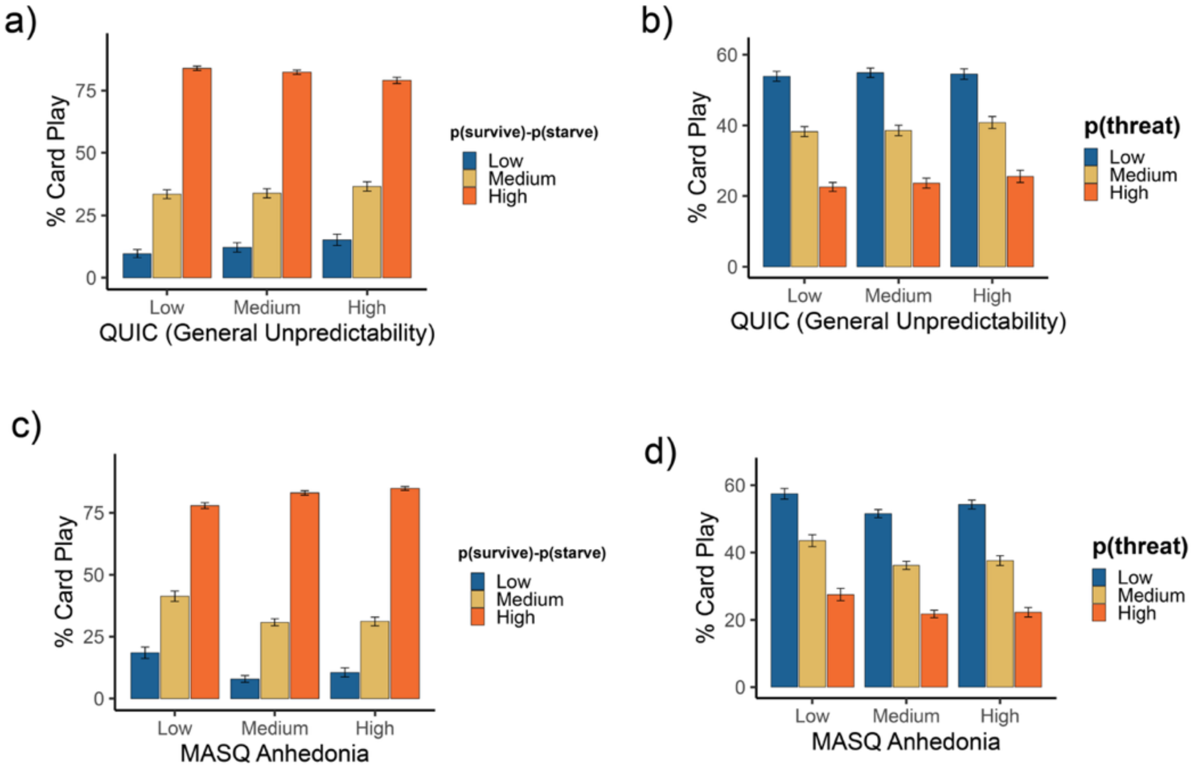


**Supplemental Figure 2. Choices by trial type across unpredictability and anhedonia tercile groups.** Trials were split into terciles based on low, medium, and high levels of motivational conflict (A & C, p(survive)-p(starve) when "Play" is chosen), as well as terciles of Zorn alien threat level (B & D). Patterns of behavior among terciles of unpredictability (A & B) and anhedonia (C & D) were then examined across the three trial types.

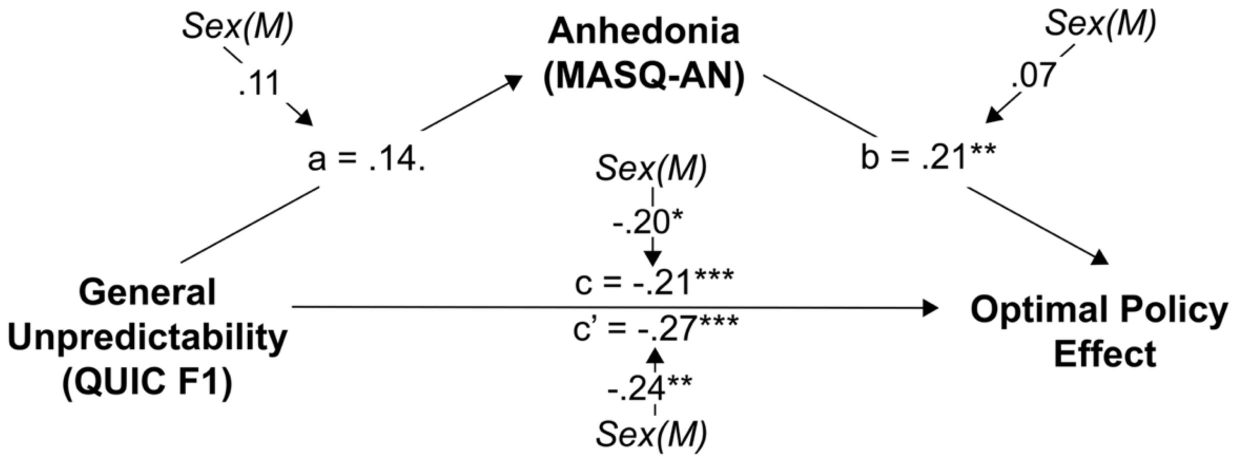


**Supplemental Figure 3. Moderated mediation analysis for sex effects.** Sex significantly moderated the relationship between unpredictability and adherence to the optimal policy in choices, where males had diminished optimal play with greater childhood unpredictability ($\beta = -.24$, $p < .001$).

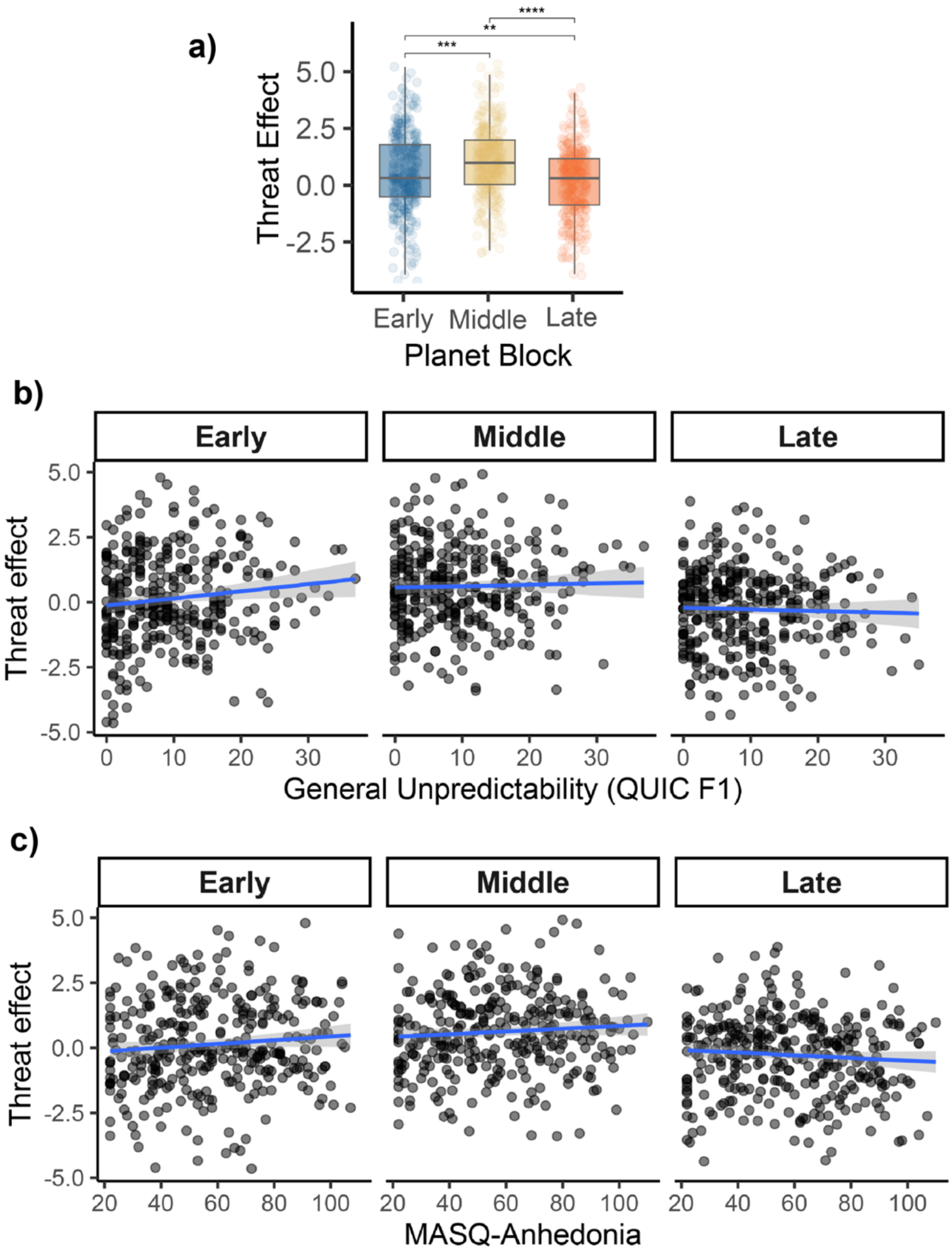


**Supplemental Figure 4. Participants' sensitivity to threat changes across the task in temporal analysis.** A) Participants were more likely to play under threat in the middle block, and more likely to wait in the late block (Kruskal-Wallis test: $\chi^2(2) = 105$, $p < .001$). B) unpredictability was associated with more significantly card play under threat in the early planets ($\beta = .027$, $t(332) = 2.27$, $p = .024$), but not in the middle ($\beta = .005$, $t(332) = .49$, $p = .62$) nor late blocks ($\beta = -.006$, $t(332) = -.59$, $p = .56$). C) Anhedonia did not significantly predict the effect of threat in any planet block (early: $\beta = .007$, $t(332) = 1.69$, $p = .09$, middle: $\beta = -.005$, $t(332) = 1.40$, $p = .16$, late: $\beta = -.005$, $t(332) = -1.42$, $p = .17$).
